# Beyond membrane permeability: A role for the small RNA MicF in regulation of chromosome replication and partitioning

**DOI:** 10.1101/2024.04.22.590647

**Authors:** Aaron Y. Stibelman, Amy Y. Sariles, Melissa K. Takahashi

## Abstract

Small regulatory RNAs (sRNA) have been shown to play a large role in the management of stress responses in *Escherichia coli* and other bacteria. sRNAs act post-transcriptionally on target mRNA through an imperfect base pairing mechanism to regulate downstream protein expression. The imperfect base pairing allows a single sRNA to bind and regulate a variety mRNA targets which can form intricate regulatory networks that connect different physiological processes for the cell’s response. Upon exposure to antimicrobials and superoxide generating agents, the MicF sRNA in *E. coli* has been shown to regulate a small set of genes involved in the management of membrane permeability. Currently, it is unknown whether MicF acts on other processes to mediate the response to these agents. Using an sRNA interaction prediction tool, we identified genes in *E. coli* that are potentially regulated by MicF. Through subsequent analysis using a sfGFP-based reporter-gene fusion, we have validated two novel targets of MicF regulation: SeqA, a negative modulator of DNA replication, and ObgE, a GTPase crucial for chromosome partitioning. Importantly, the interaction between MicF and these target mRNAs is contingent upon the presence of the RNA chaperone protein, Hfq. Furthermore, our findings affirm the role of MicF’s conserved 5’ seed pairing region in initiating these regulatory interactions. Our study suggests that, beyond its established role in membrane permeability management, MicF exerts control over chromosome dynamics in response to distinct environmental cues, implicating a more multifaceted regulatory function in bacterial stress adaptation.

## Introduction

Bacteria encounter a wide range of pressures in their natural environment such as nutrient availability, oxidative stress, and presence of antimicrobials. Small regulatory RNAs (sRNAs) aid in the adaptation to these conditions by enabling bacteria to swiftly transition between different physiological states (1, 2). These regulatory RNAs also allow bacteria to efficiently allocate resources and maximize their survival in diverse environmental niches. sRNAs modulate the expression of distally encoded *trans* mRNA by base pairing at locations near the 5’ untranslated region (UTR). The hybridization of sRNAs may alter mRNA translation directly by affecting ribosome accessibility or indirectly by modifying mRNA stability (3, 4). An important aspect of sRNA-based gene regulation is that it occurs through imperfect base-pairing interactions that allow a single sRNA to act with regulatory plasticity, targeting genes across diverse pathways, linking different biological processes, and facilitating connections within unique cellular networks (2, 5).

MicF was amongst the earliest chromosomally encoded sRNAs discovered and initially was only associated with the repression of the nonspecific outer membrane protein, OmpF (6, 7). MicF’s conservation across γ-proteobacteria (8, 9) underscored its pivotal role in responding to various extracellular stresses. MicF expression is activated by the transcription factors Rob, MarA, and SoxS, and repressed by H-NS and the leucine-responsive transcription factor, Lrp (10). In response to antibiotics or oxidative stress, among others, MicF reduces outer membrane permeability, enabling survival (11–14). Conversely, under nutrient-limiting conditions, MicF expression is suppressed to maximize OmpF production (15). More recently, additional targets of MicF regulation in *Escherichia coli* were identified including the mRNAs *lrp, cpxR*, and *phoE* (16). These genes expanded MicF’s role in membrane permeability management and suggested a larger role for MicF in metabolism via repression of Lrp which is responsible for regulation of approximately 10% of genes in *E. coli* (17).

To further explore the regulatory plasticity of MicF we used sRNA target prediction tools to identify new candidates for MicF regulation. Using mRNA-sfGFP fusions, Western blot analysis, and cell growth experiments, we identify two additional targets of MicF regulation, *obgE* and *seqA*. The repression of both genes by MicF suggests two novel roles for MicF regarding chromosome replication during conditions of nutrient abundance and oxidative stress.

## Results

### Identification of MicF regulated mRNA targets

To identify novel *E. coli* mRNA as candidates for regulation by MicF, we utilized two sRNA target prediction tools, CopraRNA (18) and TargetRNA2 (19). Both tools have previously been shown to accurately predict *in vivo* mRNA targets of bacterial sRNAs by generating RNA-RNA interactions after accounting for the intramolecular accessibility of each RNA and the phylogenetic conservation of each interaction. When evaluating the top results from both algorithms (Table S4-S5), only CopraRNA predicted known targets of MicF regulation in *E. coli* (*ompF, lrp, oppA*). Therefore, we restricted our selection of candidate mRNA targets to those exclusively predicted by the CopraRNA algorithm. We selected five of the top 10 hits (*obgE, seqA, hofQ, mgrB, hypB*) for experimental validation, by excluding those that have previously been validated (*lrp* (16), *oppA* (9), *ompF* (20)), invalidated (*murG* (16)), or had uncharacterized function (*ysaB*).

To evaluate MicF’s ability to regulate the mRNA targets predicted by CopraRNA, we built translational fusions of the mRNAs with superfolder green fluorescent protein (sfGFP) following the work of Urban and Vogel (21) and Corcoran et al. (9). When testing an sRNA’s ability to regulate an mRNA it is common to truncate the coding sequence (CDS) of the target when fusing it to the reporter protein (21, 22). As the length of the fused coding sequence could impact both sRNA binding and sfGFP protein stability, we built four variants for each target that included either 5, 10, 20, or 40 codons downstream of the predicted MicF interaction site. In the case of *mgrB*, a maximum of only 34 codons were included due to the small size of its CDS. For the predicted monocistronic targets (*seqA* and *mgrB*), the entire 5’ UTR was incorporated into the fusion. For targets predicted within an operon (*obgE, hofQ*, and *murG*), a truncated lacZ sequence was placed upstream of the target mRNA to mimic the transcription and translation of a polycistronic mRNA (21). MicF was predicted to interact with *obgE* downstream of the stop codon of the previous gene in its operon thus, the stop codon was placed at the end of the lacZ sequence, followed by the *obgE* 5’ UTR. For *hofQ* and *murG* MicF was predicted to interact with sequence within the CDS of the previous gene, thus sequence starting 30 nucleotides upstream of the predicted interaction site was fused to the lacZ sequence.

The expression of each candidate mRNA target was evaluated in the presence and absence of MicF by transforming each mRNA::sfGFP plasmid into *E. coli* BW25113 along with either a control plasmid (pControl) or one that constitutively transcribes MicF (pMicF). Of the five genes tested, *obgE::sfGFP* and *seqA::sfGFP* had variants that were repressed at least two-fold by MicF (Figure 1). Although several *mgrB::sfGFP* variants showed increased expression in the presence of MicF, we chose to only pursue targets that exhibited at least a two-fold change in expression. The *obgE* 5-codon and the *seqA* 20-codon fusions were used for all subsequent experiments.

**Figure 1.**
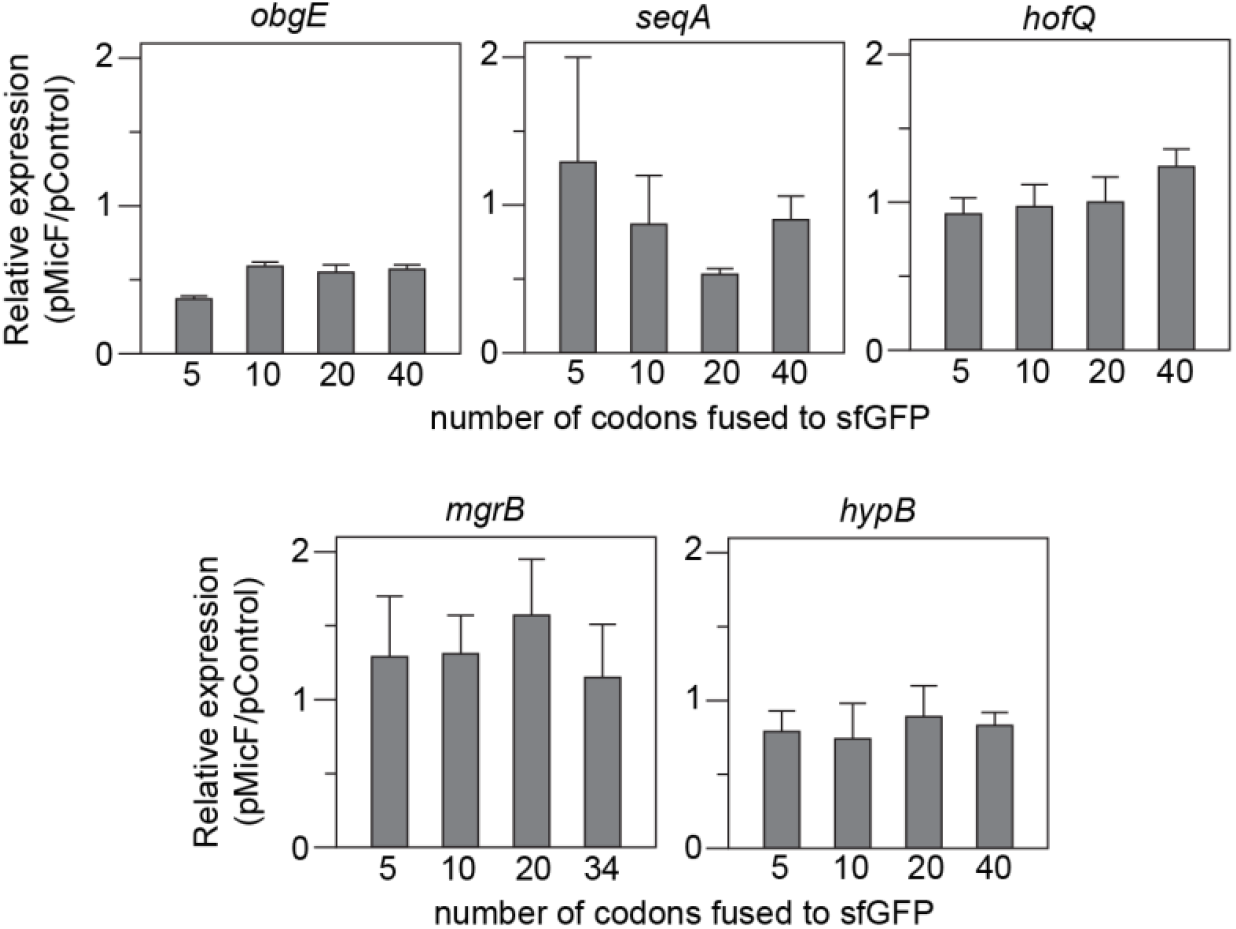
Regulation of mRNA::sfGFP reporter gene fusions by MicF. Target mRNA sequences were cloned as sfGFP fusions on a low-copy vector (pSC101) and transcribed from the constitutive promoter J23118. Four fusions were built for each target with either 5, 10, 20, or 40 codons beyond the predicted MicF binding site fused to sfGFP. mRNA::sfGFP fusions were transformed into *E. coli* BW25113 with either a MicF overexpression plasmid (pMicF) or a control (pControl), on a medium-copy vector (p15A), also transcribed from the J23118 promoter. sfGFP fluorescence and OD_600_ was measured for each condition. Each bar represents the ratio of FL/OD between cells harboring pMicF and pControl. Error bars were calculated from six biological replicates (see Figure S1).

To confirm that MicF represses the expression of ObgE and SeqA *in vivo*, we performed a Western blot analysis by chromosomally inserting a 3xFLAG epitope at the C-terminal end of either the *obgE* or *seqA* CDS and using anti-FLAG antibodies to observe protein levels in a strain of *E. coli* with the *micF* gene deleted. The production of ObgE::3xFLAG and SeqA::3xFLAG was compared between a strain complemented with MicF from either a high (pMicF ColE1) or medium (pMicF p15A) copy number plasmid and a strain transformed with a control plasmid (pControl). The Western blot analyses confirmed that MicF negatively regulates ObgE and SeqA, as their expression was decreased in the presence of MicF (Figure 2, Figure S2, Table S6).

**Figure 2.**
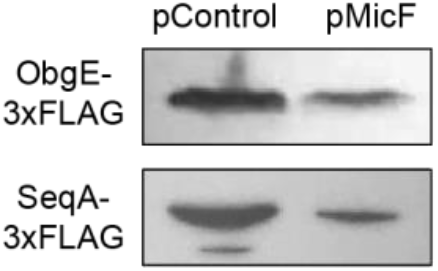
MicF represses ObgE and SeqA expression. Western blot analysis of ObgE-3xFLAG or SeqA-3xFLAG protein expressed in *E. coli* BW25113 Δ*micF* transformed with either a MicF overexpression plasmid (pMicF) or control (pControl) on a high-copy vector (ColE1). See Figure S2 and Table S6 for loading control and replicate analysis.

### The RNA chaperone protein Hfq is required for MicF’s inhibition of *obgE* and *seqA*

Due to their intrinsically weak posttranscriptional interactions with mRNA, many sRNA require the RNA chaperone protein Hfq to more stably interact with their targets (23). Hfq supports RNA-RNA interactions by binding to both molecules and causing an increase in their local concentration (24). Additionally, as Hfq binds to each RNA, it facilitates the melting of intramolecular secondary structures, further promoting sRNA-mRNA hybridization (25). Since Hfq is required for regulation of MicF’s known targets (9, 21), we investigated its necessity for the repression of *obgE* and *seqA*. This was done by transforming the mRNA::sfGFP fusions with pMicF or pControl into a Δ*hfq* strain of *E. coli* BW25113. In the Hfq deleted strain, MicF’s repression of both *obgE::sfGFP* and *seqA::sfGFP* was completely lost, suggesting that like its other targets, Hfq is required for MicF’s inhibition of *obgE* and *seqA* (Figure 3).

**Figure 3.**
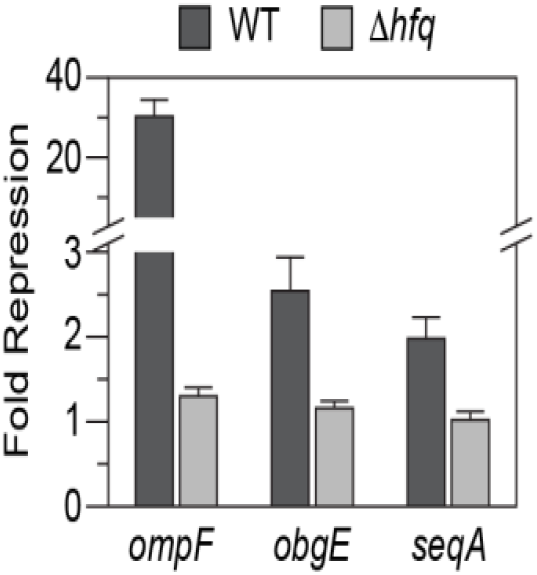
MicF’s regulation of *obgE* and *seqA* is dependent on the chaperone protein Hfq. Bars represent the ratio of FL/OD_600_ from cells with *ompF::sfGFP, obgE::sfGFP*, or *seqA::sfGFP* plasmids in the presence of a control plasmid versus one that over expresses MicF. Plasmids were transformed into an *hfq* deficient BW25113 strain of *E. coli* (Δ*hfq*) or wild type (WT). Error bars were calculated from six biological replicates (see Figure S3).

### MicF’s 13-nucleotide seed pairing region is required for the repression of *obgE* and *seqA*

A feature of many sRNAs is an unstructured region called the seed pairing region that initiates base pairing with target mRNAs (26). The seed pairing region of MicF consists of a conserved 13 nucleotides at the 5’ end that can fully regulate the mRNAs *lrp* and *ompF* when fused to an unrelated sRNA backbone (9, 27). To explore the role of MicF’s seed pairing region in the repression of *obgE* and *seqA*, we designed a synthetic sRNA composed of two parts: the first 13 nucleotides of MicF and the Hfq binding scaffold of an unrelated sRNA (SgrS) which has been used in the engineering of synthetic Hfq-dependent sRNAs (28–30) (Figure 4A-B). To test these constructs, we transformed *E. coli* BW25113 cells with one of two MicF variants. The first is composed of the seed pairing region of MicF fused to the 5’ end of the SgrS scaffold (SSM13), and the second has the first 13 nucleotides of MicF deleted (MΔ13). As expected, the repression of an *ompF::sfGFP* control was severely reduced in the absence of MicF’s seed pairing region, while its regulation by SSM13 was comparable to that of the complete MicF sequence. The repression of both *obgE::sfGFP* and *seqA::sfGFP* was eliminated in the absence of MicF’s seed pairing region. However, the seed pairing region alone was not capable of fully repressing either gene (Figure 4C).

**Figure 4.**
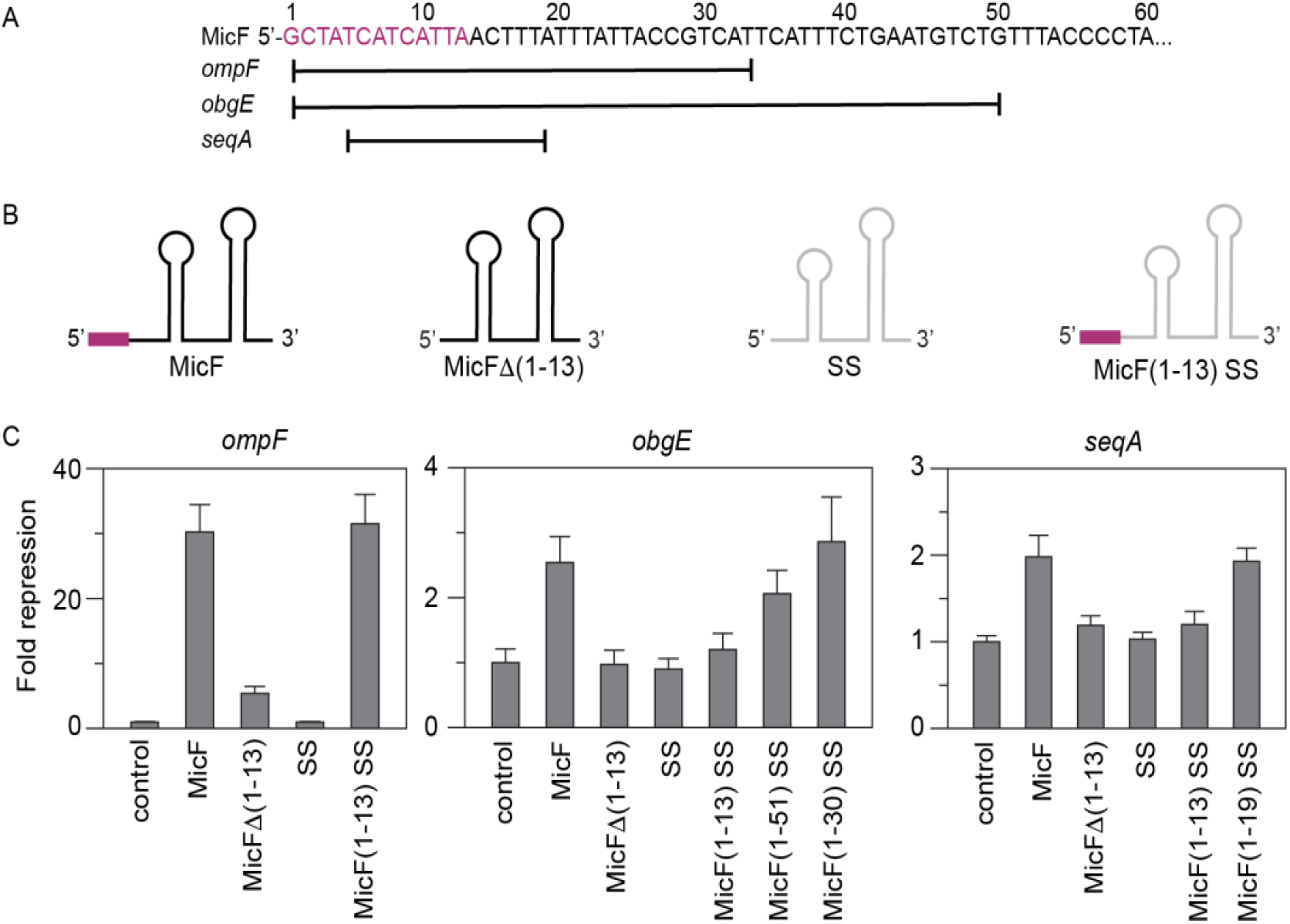
Identifying the base pair regions required for MicF’s regulation of *obgE* and *seqA*. A. The sequence for the 5’ end of MicF and its seed pairing region (in magenta). Lines indicate the nucleotides predicted to interact with the mRNAs of *ompF, obgE*, and *seqA*. A more detailed representation of the predicted interactions between *obgE* and *seqA* are displayed in Figure S4. B. Schematics of the MicF variants. The magenta box represents MicF’s 13-nucleotide seed region. SS is the SgrS scaffold used for the testing of truncated MicF segments. C. Bars represent the ratio of FL/OD_600_ from cells with *ompF::sfGFP, obgE::sfGFP*, or *seqA::sfGFP* plasmids in the presence of a control plasmid versus one that over expresses a MicF variant. Plasmids were transformed into *E. coli* BW25113. Error bars were calculated from six biological replicates (see Figure S4).

Next, we sought to identify the portion of MicF that would be sufficient for the repression of *obgE* and *seqA*. CopraRNA predicts that MicF binds to *obgE* in several different regions through nucleotide 51 of MicF while only nucleotides 5-19 are predicted to interact with *seqA* (Figure S4). We built two new constructs where nucleotides 1-51 or 1-19 were fused to the 5’ end of the SgrS scaffold, SSM51 and SSM19, respectively. While SSM19 sufficiently repressed *seqA::sfGFP*, SSM51 only moderately repressed *obgE::sfGFP* (Figure 4C). To further investigate *obgE* regulation, we considered the possibility that not all predicted interactions take place. CopraRNA does not take into account Hfq binding, which is predicted to occur somewhere within the region that spans nucleotides 28 to 93 of MicF (31). Thus, in SSM51, nucleotides 28-51 may interfere or compete with the SgrS scaffold and reduce its accessibility to Hfq. Since one way sRNAs regulate expression is through occlusion of the ribosome binding site (RBS), we used the RBS Calculator (32) to predict the RBS of *obgE* and fused the nucleotides of MicF that were predicted to bind through the RBS to the SgrS scaffold (SSM30). Using this variant, we observed complete repression of *obgE::sfGFP* (Figure 4C) suggesting that regulation of *obgE* may occur through an interference with ribosome accessibility.

### The repression of *obgE* and *seqA* is not influenced by RNase E mediated decay

While the first 19 nucleotides of MicF were sufficient to repress *seqA::sfGFP*, those nucleotides are predicted to bind within the CDS of *seqA*, downstream of both the RBS and start codon (Figure S4). This suggests an alternative mechanism for *seqA* repression that does not involve ribosome occlusion. Since sRNAs may control protein expression by modifying the stability of the mRNA transcript, we explored whether MicF exacerbates the decay of the *seqA* transcript by inducing its degradation by RNase E. In *Salmonella*, MicF represses *lpxR* through a dual mechanism, inhibition of ribosome accessibility and stimulation of degradation by RNase E (9). Therefore, we also probed MicF’s ability to induce degradation of the *obgE* transcript by RNase E.

In *E. coli*, RNase E is one of the major enzymes involved in the formation of the multi-protein RNA degradosome. The N-terminal half of RNase E contains its endoribonuclease activity while the C-terminal half is natively unstructured and serves as a scaffold for binding of the other degradosome components: PNPase, RhlB, and enolase (33). The C-terminal domain of RNase E also interacts with Hfq and this interaction is necessary for the sRNA-mediated degradation of mRNA (34). To determine whether MicF’s repression of *obgE* and *seqA* involves Hfq-dependent RNase E mediated decay, we utilized a strain of *E. coli* where the entire C-terminal scaffold of RNase E is deleted (*rne-131*) (35, 36). In the *rne-131* strain, neither *obgE::sfGFP* nor *seqA::sfGFP* was fully repressed by MicF (Figure 5).

**Figure 5.**
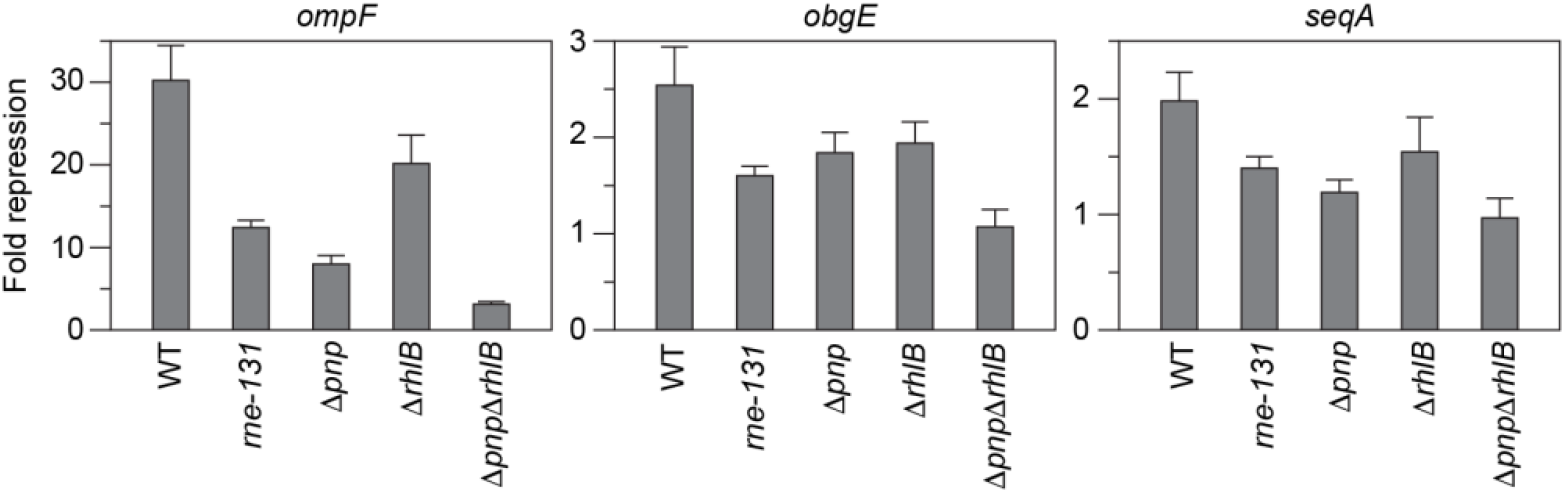
Contribution of the RNA degradosome in regulation by MicF. Bars represent the ratio of FL/OD_600_ from cells with *ompF::sfGFP, obgE::sfGFP*, or *seqA::sfGFP* plasmids in the presence of a control plasmid versus one that over expresses MicF. Plasmids were transformed into strains of *E. coli* BW25113 mutated to remove either: the C-terminal half of RNase E (*rne-131*), PNPase (Δ*pnp*), Rhlb (Δ*rhlb*), or both (Δ*pnp*Δ*rhlb*). Error bars were calculated from six biological replicates (see Figure S5).

The *rne-131* truncation also prevents PNPase and RhlB from associating with RNase E. PNPase is a 3’ exoribonuclease while RhlB is a DEAD-box RNA helicase. Both aid in the decay of RNA intermediates created by RNase E and have reduced activity when not associated to RNase E (33). Additionally a complex between PNPase, Hfq, and the sRNA protects the sRNA from degradation by RNase E for some sRNAs (37, 38). Previous work demonstrated that the regulation of *ompF* by MicF does not depend on RNase E degradation (21). In the *rne-131* strain a reduction in fold repression of *ompF::sfGFP* was observed, which suggests that the effect seen on the regulation of all three mRNAs could be due to the reduced activity of PNPase and RhlB. To assess this directly, we tested MicF regulation in strains that had an intact RNase E, but deleted PNPase (Δ*pnp*), Rhlb (Δ*rhlb*), or both (Δ*pnp*Δ*rhlb*). For all three mRNAs, we observed a reduction in fold repression in the Δ*pnp* and Δ*rhlb* strains. Furthermore, *obgE::sfGFP* and *seqA::sfGFP* were no longer repressed by MicF in the double knock out (Figure 5). Together these results suggest RNase E is not involved in MicF’s repression of *obgE* and *seqA*.

### Overexpression of MicF results in an increase in cell doubling time

In *E. coli*, the primary role of SeqA is to regulate the start of chromosome replication through its sequestration of hemimethylated origins of replication from the replication initiator protein, DnaA (39). Following replication, SeqA also aids in the cohesion of sister chromosomes prior to their segregation (40). Mutants deficient in SeqA produce filamentous cells with unsegregated DNA and their doubling times are increased by approximately 20-30% (41, 42).

To observe whether MicF overexpression would lead to phenotypes resembling *seqA* mutated strains, we observed the morphology and doubling time of Δ*micF* cells that were complemented with a plasmid that overexpresses MicF (Δ*micF* pMicF). Δ*micF* cells had comparable morphology and doubling times to that of wild type cells. Δ*micF* cells that overexpressed MicF formed filamentous cell bodies that were more variable in length and had increased doubling times when compared to wild type (Figure 6, Table 1). To determine if these observations were due to MicF’s repression of *seqA*, we simultaneously overexpressed MicF and SeqA from a plasmid in the Δ*micF* cells (Δ*micF* pMicF pSeqA). The overexpression of SeqA resulted in doubling times comparable to that of the wild type, thus linking the observed phenotype to MicF’s repression of *seqA* (Table 1).

**Figure 6.**
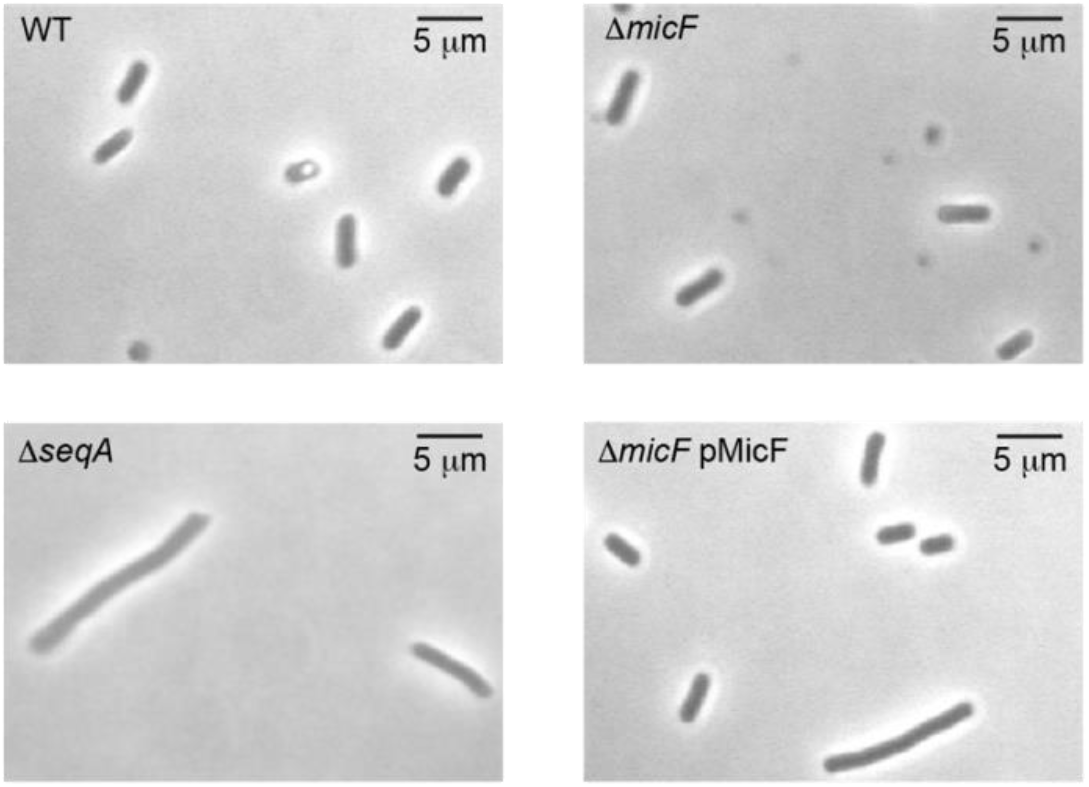
Overexpression of MicF affects cell morphology. *E. coli* BW25113 wild type, Δ*micF*, Δ*seqA*, and Δ*micF* complemented with a plasmid that overexpresses MicF (Δ*micF* pMicF) were grown to and OD_600_ of 0.2 and visualized using phase contrast. Additional observations can be found in Figure S6.

**Table 1.** Average doubling time of *E. coli* variants.

| Strain | DT (min) ± STD |
| --- | --- |
| Wild type (BW25113) | 24.0 ± 1.7 |
| $\Delta seqA$ | 32.2 ± 2.9 |
| $\Delta micF$ | 24.3 ± 1.6 |
| $\Delta micF$ pMicF | 27.6 ± 1.8 |
| $\Delta micF$ pMicF pSeqA | 25.1 ± 0.6 |

## Discussion

Here, we experimentally validated two additional mRNA targets of MicF (*obgE* and *seqA*) that were predicted by CopraRNA. ObgE is an essential GTPase in *E. coli* that has roles in chromosome partitioning (43, 44) and in the cellular response to amino acid starvation (45). SeqA serves as a negative modulator during the initiation of chromosome replication (39) that also facilitates the segregation of newly replicated sister chromosomes (40). We found that MicF represses expression of both genes.

Our experiments confirmed that MicF regulates both genes through an Hfq-dependent antisense mechanism that requires the 5’ seed pairing region of MicF. Unlike the regulation of OmpF and Lrp (9), the seed pairing region alone was not sufficient to repress *obgE::sfGFP* and *seqA::sfGFP* (Figure 4). The regulation of ObgE included the first 30 nucleotides of MicF that are predicted to bind through the RBS and start codon of *obgE*. The seed region itself is not involved in the pairing with the RBS or start codon (Figure S4), which suggests its role is to initiate binding between the two RNAs. Part of MicF’s seed pairing region is predicted to bind to *seqA* although the first 19 nucleotides were required to completely repress *seqA::sfGFP*. In the case of SeqA, MicF is only predicted to bind within the CDS and not the RBS or start codon. It is still possible that MicF inhibits translation initiation even though it does not bind to the RBS or start codon. Previous work investigating RybB repression of *ompN* in *Salmonella* demonstrated that translational control could occur if base pairing took place within a five codon window (46). MicF is predicted to bind to the *seqA* CDS starting within the fourth codon.

Besides occluding translation initiation, sRNAs are known to repress protein expression by promoting mRNA decay by RNase E or RNase III (47, 48). RNase III is known to degrade dsRNA or intramolecular duplexes formed within a single RNA, however, in either case the duplex must be of sufficient length, approximately 22 base pairs (49). In the case of *seqA*, the predicted binding interaction is only 15 base pairs long, which makes it a less likely target. We assessed the role of RNase E, however, neither the regulation of SeqA nor ObgE appeared to be dependent on RNase E (Figure 5). Given these results, we propose that MicF represses translation of both ObgE and SeqA by preventing formation of the translation initiation complex.

The regulation of ObgE and SeqA introduces a new role for MicF beyond regulation of the outer membrane. ObgE is known to play a critical role in chromosome partitioning and is potentially used in a replication checkpoint response (44, 50). Notably, chromosome partitioning defects have been observed with a modest reduction in ObgE concentrations (43). Thus, repression of ObgE by MicF could contribute to cell cycle arrest. This helps to explain a role of MicF when activated by SoxS. Transcription of *soxS* is activated by SoxR, which is induced by superoxide generating agents or nitric oxide (51). Early work showed that MicF repressed OmpF due to the activation of the *soxRS* locus although the importance of OmpF suppression under oxidative stress conditions was not clear (13). Strains deficient in *soxRS* were shown to be hypersensitive to chloramphenicol and nalidixic acid which does suggest a direct role for repression of OmpF since one way these antibiotics enter the cell is through OmpF (52, 53). However, elimination of OmpF did not increase the resistance of *E. coli* to menadione, another agent known to induce *soxRS* (13). Other work showed only a modest increase in Lrp expression in a Δ*micF* strain that was treated with paraquat (54). Perhaps the role of MicF during oxidative stress is to suppress chromosome partitioning and work together with the OxyS sRNA to induce cell cycle arrest and provide time for DNA damage repair (55)

SeqA has also been tied to replication arrest, however, this is under conditions with high SeqA concentrations (56, 57). Instead, we propose a role for suppression of SeqA under high nutrient conditions where MicF is already known to repress OmpF and Lrp (10, 16). Initiation of chromosome replication by DnaA is blocked by binding of SeqA to hemimethylated *oriC*. This prevents immediate re-initiation and ensures a single round of replication per division cycle. However, under nutrient-rich conditions and faster growth rates it is known that *E. coli* can initiate a new round of replication before the previous round is complete (39, 58). DnaA concentration increases with increasing growth rate (59) and a parallel decrease in SeqA concentration would aid in initiation of another round of replication. Repression of SeqA under these conditions would complement MicF’s repression of OmpF and Lrp.

### Experimental Procedures

#### Growth conditions

All strains were grown in Lysogeny broth (LB), LB agar (1.5%) plates, or MOPS EZ Rich Defined Media (Teknova M2105) at 37°C. When necessary, media was supplemented with antibiotics (carbenicillin: 100 μg/ml, chloramphenicol: 34 μg/ml, kanamycin: 50 μg/ml).

#### Strains

The *E. coli* strains used in this study are listed in Table S1. Deletion strains were constructed using the lambda red recombination method (60). DNA fragments containing a KanR cassette flanked by two FRT sites were amplified from pKD4 with the appropriate flanking sequences and electroporated into BW25113 for the construction of Δ*micF* and rne-131, or into a Keio strain (Δ*rhlB*) (61) for the construction of Δ*rhlB*Δ*pnp*. The construction of the C-terminal 3xFLAG tagged strains were carried out similarly using the following modification. The DNA fragments contained a 3xFLAG tag followed by a FRT flanked CmR cassette. DNA fragments were PCR amplified and electroporated into the Δ*micF* strain. Removal of KanR or CmR was achieved using the helper plasmid pCP20 following Datsenko and Wanner (60). All deletions and insertions were confirmed using PCR and Sanger DNA sequencing. Primers used for creating the deletion strains or 3xFLAG tag strains are listed in Table S1.

#### Plasmids

All plasmids used in this study are listed in Table S3, with key sequences found in Table S2. Plasmids were constructed using inverse PCR or Gibson assembly. Gene sequences were PCR amplified from purified genomic DNA. New England Biolabs Turbo *E. coli* cells was used for transformation of constructed plasmids. All plasmids were verified using Sanger DNA sequencing.

#### CopraRNA and TargetRNA2

The web-accessible programs CopraRNA and TargetRNA2 were used to identify target genes regulated by MicF in *E. coli*. The organisms included in the CopraRNA query were *E. coli* NC 000913, *Citrobacter koseri* NC009792, *Citrobacter rodentium* NC_013716, *Escherichia fergusonii*, NC 011740, and *Salmonella* enterica subsp. enterica serovar *Typhimurium* NC 003197. The CopraRNA program was run under default conditions. The CopraRNA default consists of sequences extracted around the start codon (200 nucleotides upstream, 100 nucleotides downstream), a dynamic setting for p-value, and no p-value filtering or consensus prediction. For the TargetRNA2, the MicF sequence was obtained from Genbank NC_000913 and screened against the genome of E. coli str. K-12 substr. MG1655.

#### Fluorescence measurement and culturing conditions

Plasmids were transformed into chemically competent *E. coli* strains, plated on LB-agar plates containing chloramphenicol and carbenicillin, and incubated overnight at 37°C. Three colonies from each condition were inoculated into 300 μl of LB with corresponding antibiotics in a 2 ml 96-well block (Costar 3961) sealed with a breath-easier membrane and grown for 17 hours overnight at 37°C while shaking at 100 rpm (Labnet Vortemp 56). Four microliters of the overnight culture were added to 296 μl of MOPS media that was pre-warmed at 37°C for 30 minutes. The cultures were grown under the same conditions as above for 2.5 or 3 hours depending on the strain used. One hundred microliters of each culture was transferred to a 96-well plate (Costar 3631), and OD600 and bulk sfGFP fluorescence (485 nm excitation, 520 nm emission) were measured using a Biotek Synergy H1 plate reader. Each experiment included two sets of controls: a media blank and *E. coli* transformed with control plasmids. OD and FL for each culture were first corrected by subtracting the mean value of the media blank. The ratio of the corrected FL and OD (FL/OD) was calculated for each culture. The *E. coli* cultures transformed with control plasmids were used to correct for autofluorescence. The average FL/OD from the control cultures was subtracted from FL/OD from each condition.

#### Determining cell doubling time

*E. coli* strains were isolated on LB-agar plates containing appropriate antibiotics if necessary. Six individual colonies from each condition were inoculated into 300 μl of LB with corresponding antibiotics, if necessary, in a 2 ml 96-well block (Costar 3961) sealed with a breath-easier membrane and grown for 17 hours overnight at 37°C while shaking at 100 rpm (Labnet Vortemp 56). Four microliters of the overnight culture were added to 296 μl of LB and grown under the same conditions for one hour. From that point, 2 μl from each culture was removed every 20 minutes and used to determine CFU/ml via dilutions and plating. The doubling time for an individual colony was determined by taking the log base 2 of each CFU/ml value, then determining the slope log_2_(CFU/ml) vs time and taking the inverse of that slope value. Doubling times were averaged across the six colonies.

#### Protein purification and Western blot

*E. coli* cultures were grown overnight at 37°C while shaking at 275 rpm in LB supplemented with appropriate antibiotics. Cells were diluted to an OD600 of 0.02 in LB and grown to an OD600 of 0.5. Cell pellets corresponding to 1 ml of each culture were resuspended in 2x loading buffer and 2M DTT, and heated for 5 minutes at 95°C. The 2x loading buffer contains 10% SDS, 50% glycerol, 1M Tris-HCl (pH 6.8), and 0.2 mg/ml Bromophenol Blue. For each sample, aliquots of 10 μl were separated on either a 10% or 15% SDS-PAGE gel and transferred to a 0.2 μM PVDF membrane (Sigma Aldrich 0301004001) at 10 mA overnight at 4°C, with stirring. Protein expression was visualized using monoclonal anti-FLAG antibodies, HRP (Thermo Fisher MA1-91878-HRP), and developed using Pierce Fast Western Kit SuperSignal West Pico Mouse (Thermo Fisher 35060). Blots were developed in SuperSignal West Pico Working Solution for five minutes and 2-minute exposed films (Thomas scientific 1148B77) were developed on an SRX 101A X-ray film processor (Konica Minolta). The images were processed and quantified using the Image J software. The intensities of the MicF target protein bands were normalized with the intensities of the corresponding total protein visualized via Ponceau S staining. For the normalization, the intensities of the MicF target protein bands were divided by that of the total protein intensity.

#### Observation of cell growth via phase contrast microscopy

*E. coli* strains transformed with a control plasmid (pMKT046) or MicF expression plasmid (pMKT050) were isolated onto LB agar supplemented with kanamycin. A single colony from each strain was grown overnight at 37°C while shaking at 275 rpm in LB supplemented with kanamycin. The next day, cultures were diluted 100-fold into LB and grown until they reached an OD600 of 0.2. A wet mount was prepared, and morphology was visualized via phase contrast microscopy using a Nikon E200 microscope at 400x.

## Data Availability

Data are contained within the manuscript and Supporting Information. Additional raw data is available upon request from the corresponding author.

## Supporting Information

This article contains supporting information: Tables S1-S6, and Figures S1-S7. Reference 60 is repeated in the supporting information.

## Acknowledgements

We would like to thank Cristian Ruiz Rueda, Gilberto Flores, and Mariano Loza Coll (California State University Northridge) for experimental protocols and constructive feedback on this manuscript.

## Author contributions

All authors performed experiments, analyzed data, and wrote the manuscript.

## Funding and additional information

Funding for this research was provided by California State University Northridge start-up funds to M.K.T. Research reported in this publication was supported by the National Institute of General Medical Sciences of the National Institutes of Health under Award Number TL4GM118977 to A.Y.S.-2. The content is solely the responsibility of the authors and does not necessarily represent the official views of the National Institutes of Health.

