## Supplemental Information for "Beyond membrane permeability: A role for the small RNA MicF in regulation of chromosome replication and partitioning"

Table of contents:


| | **Description** | **Page** |
|---|---|---|
| Table S1 | Strains used in this study | 2 |
| Table S2 | Important DNA sequences | 3-5 |
| Table S3 | Plasmids used in this study | 6 |
| Table S4 | Top 15 CopraRNA results | 7 |
| Table S5 | Top 15 TargetRNA2 results | 7 |
| Figure S1 | FL/$OD_{600}$ measurements for data in Figure 1 | 8 |
| Figure S2 | Controls and additional replicates for Western blots in Figure 2 | 9 |
| Table S6 | Image J analysis of band intensities for ObgE-3xFLAG | 9 |
| Figure S3 | FL/$OD_{600}$ measurements for data in Figure 3 | 10 |
| Figure S4 | FL/$OD_{600}$ measurements for data in Figure 4 | 11 |
| Figure S5 | FL/$OD_{600}$ measurements for data in Figure 5 | 12 |
| Figure S6 | Additional observations of cell morphology (supplemental to Figure 6) | 13 |
| Figure S7 | Doubling time measurements for data in Table 1 | 13 |
| References | | 13 |

**Table S1**: Strains used in this study. Includes primers used to create strains made for this study.

| Strain | Reference | Forward primer | Reverse primer |
|---|---|---|---|
| *E. coli* BW25113 | | | |
| *E. coli* BW25113 Δ*hfq* | (1) | | |
| *E. coli* BW25113 Δ*rhlB* | (1) | | |
| *E. coli* BW25113 Δ*pnp* | (1) | | |
| *E. coli* BW25113 Δ*micF* | This study | GTCAAAACAAACCTTCACTCGCAACTAGAATAACTCCCGGTGTAGGCTGGAGCTGCTTC | AGTGTGTAAAGAAGGGTAAAAAAAACCGAATGCGAGGCATATGGGAATTAGCCATGGTCC |
| *E. coli* BW25113 *rne-131* | This study | CTTCTTCGGCGCACTGAAAGCGCTGTTCAGCGGTGGTTAAGTGTAGGCTGGAGCTGCTTC | GATTACTTTGAGCTAATTATTACTCAACAGGTTGCGGACGATGGGAATTAGCCATGGTCC |
| *E. coli* BW25113 Δ*rhlB* Δ*pnp* | This study | CGTAAGGTACTGTCTAAGAAAGAGAAAGGATATTACATTGGTGTAGGCTGGAGCTGCTTC | CGGAGGGCAAATGGCAACCTTACTCGCCCTGTTCAGCAGCATGGGAATTAGCCATGGTCC |
| *E. coli* BW25113 Δ*micF* *obgE*:3xFLAG | This study | CGAAGACGACGAAGAAGGCGTTGAGTTCATTTACAAGCGTGACTACAAAGACCATGACGG | AAATCGTGCAAATTCAACATATTGCAATTCTCTTGTAGGCATGGGAATTAGCCATGGTCC |
| *E. coli* BW25113 Δ*micF* *seqA*:3xFLAG | This study | ATTCCCGGCGGAATTGATTGAGAAGGTTTGCGGAACTATCGACTACAAAGACCATGACGG | TGGGCGACGTTAATCAAATCACTCTGTTGTGCAGGTTGGCATGGGAATTAGCCATGGTCC |

**Table S2:** Important DNA sequences

| **Name** | **Sequence** |
|---|---|
| J23118 promoter | TTGACGGCTAGCTCAGTCCTAGGTATTGTGCTAGC |
| J23119 promoter | TTGACAGCTAGCTCAGTCCTAGGTATAATACTAGT |
| Plux promoter | ACCTGTAGGATCGTACAGGTTTACGCAAGAAAATGGTTTGTTAT<br>AGTCGAATAAA |
| LuxR | ATGAAAAACATAAATGCCGACGACACATACAGAATAATTAATAA<br>AATTAAAGCTTGTAGAAGCAATAATGATATTAATCAATGCTTATC<br>TGATATGACTAAAATGGTACATTGTGAATATTATTTACTCGCGAT<br>CATTTATCCTCATTCTATGGTTAAATCTGATATTTCAATCCTAGA<br>TAATTACCCTAAAAAATGGAGGCAATATTATGATGACGCTAATTT<br>AATAAAATATGATCCTATAGTAGATTATTCTAACTCCAATCATTC<br>ACCAATTAATTGGAATATATTTGAAAACAATGCTGTAAATAAAAA<br>ATCTCCAAATGTAATTAAAGAAGCGAAAACATCAGGTCTTATCA<br>CTGGGTTTAGTTTCCCTATTCATACGGCTAACAATGGCTTCGGA<br>ATGCTTAGTTTTGCACATTCAGAAAAAGACAACTATATAGATAGT<br>TTATTTTTACATGCGTGTATGAACATACCATTAATTGTTCCTTCT<br>CTAGTTGATAATTATCGAAAAATAAATATAGCAAATAATAAATCA<br>AACAACGATTTAACCAAAAGAGAAAAAGAATGTTTAGCGTGGGC<br>ATGCGAAGGAAAAAGCTCTTGGGATATTTCAAAAATATTAGGTT<br>GCAGTGAGCGTACTGTCACTTTCCATTTAACCAATGCGCAAATG<br>AAACTCAATACAACAAACCGCTGCCAAAGTATTTCTAAAGCAAT<br>TTTAACAGGAGCAATTGATTGCCCATACTTTAAAAATTAA |
| T1 terminator | GCATCAAATAAAACGAAAGGCTCAGTCGAAAGACTGGGCCTTT<br>CGTTTTATCTGTTGTTTGTCGGTGAACGCTCTCCTGAGTAGGAC<br>AAATCCGCCGCCCTAGA |
| MicF | GCTATCATCATTAACTTTATTTATTACCGTCATTCATTTCTGAATG<br>TCTGTTTACCCCTATTTCAACCGGATGCCTCGCATTCGGTTTTTT<br>TT |
| SeqA CDS | ATGAAAACGATTGAAGTTGATGATGAACTCTACAGCTATATTGC<br>CAGCCACACTAAGCATATCGGCGAGAGCGCATCCGACATTTTA<br>CGGCGTATGTTGAAATTTTCCGCCGCATCACAGCCTGCTGCTC<br>CGGTGACGAAAGAGGTTCGCGTTGCGTCACCTGCTATCGTCGA<br>AGCGAAGCCGGTCAAAACGATTAAAGACAAGGTTCGCGCAATG<br>CGTGAACTTCTGCTTTCGGATGAATACGCAGAGCAAAAGCGAG<br>CGGTCAATCGCTTTATGCTGCTGTTGTCTACACTATATTCTCTTG<br>ACGCCCAGGCGTTTGCCGAAGCAACGGAATCGTTGCACGGTC<br>GTACACGCGTTTACTTTGCGGCAGATGAACAAACGCTGCTGAA<br>AAATGGTAATCAGACCAAGCCGAAACATGTGCCAGGCACGCCG<br>TATTGGGTGATCACCAACACCAACACCGGCCGTAAATGCAGCA<br>TGATCGAACACATCATGCAGTCGATGCAATTCCCGGCGGAATT<br>GATTGAGAAGGTTTGCGGAACTATCTAA |
| sfGFP<br>(no start codon) | AGCAAAGGAGAAGAACTTTTCACTGGAGTTGTCCCAATTCTTGT<br>TGAATTAGATGGTGATGTTAATGGGCACAAATTTTCTGTCCGTG<br>GAGAGGGTGAAGGTGATGCTACAAACGGAAAACTCACCCTTAA<br>ATTTATTTGCACTACTGGAAAACTACCTGTTCCGTGGCCAACAC<br>TTGTCACTACTCTGACCTATGGTGTTCAATGCTTTTCCCGTTATC<br>CGGATCACATGAAACGGCATGACTTTTTCAAGAGTGCCATGCC<br>CGAAGGTTATGTACAGGAACGCACTATATCTTTCAAAGATGACG<br>GGACCTACAAGACGCGTGCTGAAGTCAAGTTTGAAGGTGATAC |

| | |
|---|---|
| | CCTTGTTAATCGTATCGAGTTAAAGGGTATTGATTTTAAAGAAGA<br>TGGAAACATTCTTGGACACAAACTCGAGTACAACTTTAACTCAC<br>ACAATGTATACATCACGGCAGACAAACAAAAGAATGGAATCAAA<br>GCTAACTTCAAAATTCGCCACAACGTTGAAGATGGTTCCGTTCA<br>ACTAGCAGACCATTATCAACAAAATACTCCAATTGGCGATGGCC<br>CTGTCCTTTTACCAGACAACCATTACCTGTCGACACAATCTGTC<br>CTTTCGAAAGATCCCAACGAAAAGCGTGACCACATGGTCCTTCT<br>TGAGTTTGTAACTGCTGCTGGGATTACACATGGCATGGATGAG<br>CTCTACAAA |
| *obgE* (stop codon of previous gene through 40 codons) | GCATACAACGGTGGTATCGCAACCCCGCGCAGGCGAATGATTT<br>ACGGAGAATAAAATGAAGTTTGTTGATGAAGCATCGATTCTGGT<br>CGTTGCAGGTGATGGCGGTAATGGTTGCGTGAGCTTCCGCCG<br>CGAAAAGTATATTCCGAAAGGCGGCCCGGATGGCGGCGACGG<br>CGGTGATGGTGGTGACGTATGGATG |
| *seqA* (+1 through 40 codons) | ACTCCTGGCGACTTGTATTCAGCTAAGACACTGCACTGGATTAA<br>GATGAAAACGATTGAAGTTGATGATGAACTCTACAGCTATATTG<br>CCAGCCACACTAAGCATATCGGCGAGAGCGCATCCGACATTTT<br>ACGGCGTATGTTGAAATTTTCCGCCGCATCACAG |
| *hofQ* (30nt upstream of interaciton site through 40 codons) | ACTTAGCCAGTGGCGCTATCAGGGGATGGTAGGGCGAGGCGA<br>GCGCATCATCGGTGTAATAAAAGACGGGCAAAAGAAATGGCGA<br>CGGGTGCAGCAAAACGATGTGCTGGAAAACGGCTGGACAATTT<br>TACAGCTGACGCCAGACGTACTAACGCTGGGTACCGGGACAAA<br>CTGCGAACCGCCACAATGGTTGTGGCAACGGCAAGGAGATACA<br>AATGAAGCAATGGATAGCCGCACTACTGTTGATGCTGATACCC<br>GGCGTACAGGCGGCAAAGCCGCAAAAAGTGACGCTGATGGTG<br>GATGACGTTCCGGTAGCTCAGGTGTTGCAGGCGCTG |
| *mgrB* (+1 through 34 codons beyond interaction site) | ATAAGGTAGGTGAAACGGAGATTGGAGTGAAAAAGTTTCGATG<br>GGTCGTTCTGGTTGTCGTGGTGTTGGCTTGCTTGCTGCTTTGG<br>GCGCAGGTATTCAACATGATGTGCGATCAGGATGTACAATTTTT<br>CAGCGGAATTTGTGCCATTAACCAGTTTATCCCGTGG |
| *hybB* (+1 through 40 codons) | GAAGAACAAGAGGCCGAATGCTGGTGTGAAACATGCCAACAGT<br>ATGTGACGCTACTGACCCAGCGCGTCCGCCGCTGTCCACAGTG<br>TCATGGTGACATGCTGCAGATTGTGGCAGACGACGGTTTACAG<br>ATTCGGCGGATAGAAATAGACCAGGAGTGAGCGATGTGTACAA<br>CATGCGGTTGCGGTGAAGGCAACCTGTATATCGAGGGTGATGA<br>ACATAACCCTCATTCCGCGTTTCGTAGCGCGCCATTTGCCCCG<br>GCGGCACGCCCGAAGATGAAAATC |
| RBS-FlacZ | GAATTCATTAAAGAGGAGAAAGGTACCATGGACTACAAAGACCA<br>TGACGGTGATTATAAAGATCATGATATCGACTACAAAGATGACG<br>ACGATAAAACCATGATTACGGATTCACTGGCCGTCGTTTTACAA<br>CGTCGTGACTGGGAAAACCCTGGCGTTACCCAACTTAATCGCC<br>TTGCAGCACATCCCCCTTTCGCCAGCTGGCGTAATAGCGAAGA<br>GGCCCGCACCGATCGCCCTTCCCAACAGTTGCGCAGCCTGAAT<br>GGCGAATGGATGCAT |
| SgrS scaffold (SS) | TATTGGTGTAAAATCACCCGCCAGCAGATTATACCTGCTGGTTT<br>TTTTT |
| MicF(1-13)-SS | GCTATCATCATTATATTGGTGTAAAATCACCCGCCAGCAGATTA<br>TACCTGCTGGTTTTTTTT |
| MicF(1-19)-SS | GCTATCATCATTAACTTTATATTGGTGTAAAATCACCCGCCAGC<br>AGATTATACCTGCTGGTTTTTTTT |

| | |
|---|---|
| MicF(1-30)-SS | GCTATCATCATTAACTTTATTTATTACCGTTATTGGTGTAAAATC ACCCGCCAGCAGATTATACCTGCTGGTTTTTTTT |
| MicF(1-51)-SS | GCTATCATCATTAACTTTATTTATTACCGTCATTCATTTCTGAATG TCTGTTATTGGTGTAAAATCACCCGCCAGCAGATTATACCTGCT GGTTTTTTTT |
| 3xFLAG | GACTACAAAGACCATGACGGTGATTATAAAGATCATGATATCGA CTACAAAGATGACGACGATAAA |

**Table S3**: Plasmids used in this study

| Plasmid | Description | Origin/Resistance | Figure |
|---|---|---|---|
| MKT176 | J23118-T1 | p15A/AmpR | 1, 3, 4, 5, S1, S2, S3, S4 |
| MKT173 | J23118-MicF-T1 | p15A/AmpR | 1, 3, 4, 5, S1, S3, S4 |
| MKT221 | J23118-T1 | pSC101/CmR | 1, 3, 4, 5, S1, S3, S4 |
| MKT602 | J23118-RBS-FlacZ-obgE(5 codons)-sfgfp-T1 | pSC101/CmR | 1, 3, 4, 5, S1, S3, S4 |
| MKT603 | J23118-RBS-FlacZ-obgE(10 codons)-sfgfp-T1 | pSC101/CmR | 1, S1 |
| MKT604 | J23118-RBS-FlacZ-obgE(20 codons)-sfgfp-T1 | pSC101/CmR | 1, S1 |
| MKT605 | J23118-RBS-FlacZ-obgE(40 codons)-sfgfp-T1 | pSC101/CmR | 1, S1 |
| MKT419 | J23118-seqA(5 codons)-sfgfp-T1 | pSC101/CmR | 1, S1 |
| MKT420 | J23118-seqA(10 codons)-sfgfp-T1 | pSC101/CmR | 1, S1 |
| MKT421 | J23118-seqA(20 codons)-sfgfp-T1 | pSC101/CmR | 1, 3, 4, 5, S1, S3, S4 |
| MKT437 | J23118-seqA(40 codons)-sfgfp-T1 | pSC101/CmR | 1, S1 |
| MKT524 | J23118-RBS-FlacZ-hofQ(5 codons)-sfgfp-T1 | pSC101/CmR | 1, S1 |
| MKT525 | J23118-RBS-FlacZ-hofQ(10 codons)-sfgfp-T1 | pSC101/CmR | 1, S1 |
| MKT526 | J23118-RBS-FlacZ-hofQ(20 codons)-sfgfp-T1 | pSC101/CmR | 1, S1 |
| MKT539 | J23118-RBS-FlacZ-hofQ(40 codons)-sfgfp-T1 | pSC101/CmR | 1, S1 |
| MKT401 | J23118-mgrB(5 codons)-sfgfp-T1 | pSC101/CmR | 1, S1 |
| MKT402 | J23118-mgrB(10 codons)-sfgfp-T1 | pSC101/CmR | 1, S1 |
| MKT403 | J23118-mgrB(20 codons)-sfgfp-T1 | pSC101/CmR | 1, S1 |
| MKT427 | J23118-mgrB(34 codons)-sfgfp-T1 | pSC101/CmR | 1, S1 |
| MKT447 | J23118-hypB(5 codons)-sfgfp-T1 | pSC101/CmR | 1, S1 |
| MKT425 | J23118-hypB(10 codons)-sfgfp-T1 | pSC101/CmR | 1, S1 |
| MKT426 | J23118-hypB(20 codons)-sfgfp-T1 | pSC101/CmR | 1, S1 |
| MKT448 | J23118-hypB(40 codons)-sfgfp-T1 | pSC101/CmR | 1, S1 |
| MKT433 | J23118-MicF∆(1-13)-T1 | p15A/AmpR | 4 |
| MKT431 | J23118-SgrS Scaffold (SS)-T1 | p15A/AmpR | 4 |
| MKT432 | J23118-MicF(1-13)-SS-T1 | p15A/AmpR | 4 |
| MKT517 | J23118-MicF(1-19)-SS-T1 | p15A/AmpR | 4 |
| MKT572 | J23118-MicF(1-30)-SS-T1 | p15A/AmpR | 4 |
| MKT511 | J23118-MicF(1-51)-SS-T1 | p15A/AmpR | 4 |
| MKT046 | J23119-T1 | ColE1/KanR | 2, S2 |
| MKT050 | J23119-MicF-T1 | ColE1/KanR | 2, 6, T1, S2, S5, S6, S7 |
| MKT161 | J23119-MicF-T1 | p15A/AmpR | S2 |
| MKT628 | J23119-MicF-T1-PluxR-seqA-T1 | ColE1/KanR | T1, S7 |

**Table S4**: Top 15 CopraRNA results. Previously validated targets are highlighted in yellow.

| Rank | Gene Name | Gene Annotation | MicF predicted nucleotides |
|---|---|---|---|
| 1 | *lrp* | leucine-responsive global transcriptional regulator | 1 - 64 |
| 2 | *oppA* | oligopeptide ABC transporter periplasmic binding protein | 1 - 59 |
| 3 | *murG* | N-acetylglucosaminyl transferase | 1 - 51 |
| 4 | *ompF* | outer membrane porin 1a (Ia;b;F) | 1 - 33 |
| 5 | *ysaB* | uncharacterized protein | 3 - 54 |
| 6 | *obgE* | GTPase involved in cell partitioning and DNA repair | 1 - 51 |
| 7 | *seqA* | negative modulator of initiation of replication | 5 - 19 |
| 8 | *hofQ* | DNA catabolic putative fimbrial transporter | 27 - 88 |
| 9 | *mgrB* | regulatory peptide for PhoPQ feedback inhibition | 1 - 63 |
| 10 | *hypB* | GTP hydrolase involved in nickel liganding into hydrogenases | 1 - 53 |
| 11 | *mobA* | molybdopterin-guanine dinucleotide synthase | 15 - 59 |
| 12 | *smrB* | putative DNA endonuclease | 16 - 57 |
| 13 | *wza* | colanic acid export protein; outer membrane auxillary lipoprotein | 5 - 27 |
| 14 | *hsdS* | specificity determinant for hsdM and hsdR | 27 - 58 |
| 15 | *gltL* | glutamate/aspartate ABC transporter ATPase | 1 - 25 |

**Table S5**: Top 15 TargetRNA2 results

| Rank | Gene Name | Gene Annotation | MicF predicted nucleotides |
|---|---|---|---|
| 1 | *sbcB* | exodeoxyribonuclease I; exonuclease I | 23 - 39 |
| 2 | *ppsA* | phosphoenolpyruvate synthase | 59 - 74 |
| 3 | *ylcG* | uncharacterized protein, DLP12 prophage | 27 - 44 |
| 4 | *metL* | Bifunctional aspartokinase/homoserine dehydrogenase 2 | 46 - 60 |
| 5 | *ybaY* | outer membrane lipoprotein | 29 - 42 |
| 6 | *serA* | D-3-phosphoglycerate dehydrogenase | 44 - 58 |
| 7 | *nimR* | putative DNA-binding transcriptional regulator | 61 - 75 |
| 8 | *appB* | cytochrome bd-II oxidase, subunit II | 45 - 60 |
| 9 | *gltD* | glutamate synthase, 4Fe-4S protein, small subunit | 46 - 60 |
| 10 | *rmsD* | 16S rRNA m(2)G966 methyltransferase, SAM-dependent | 77 - 93 |
| 11 | *gspD* | general secretory pathway component, cryptic | 28 - 43 |
| 12 | *ebgR* | transcriptional repressor | -8 - 7 |
| 13 | *murI* | glutamate racemase | 65 - 75 |
| 14 | *ydjZ* | TVP38/TMEM64 family inner membrane protein | 1 - 20 |
| 15 | *lsrK* | autoinducer-2 (AI-2) kinase | 44 - 54 |

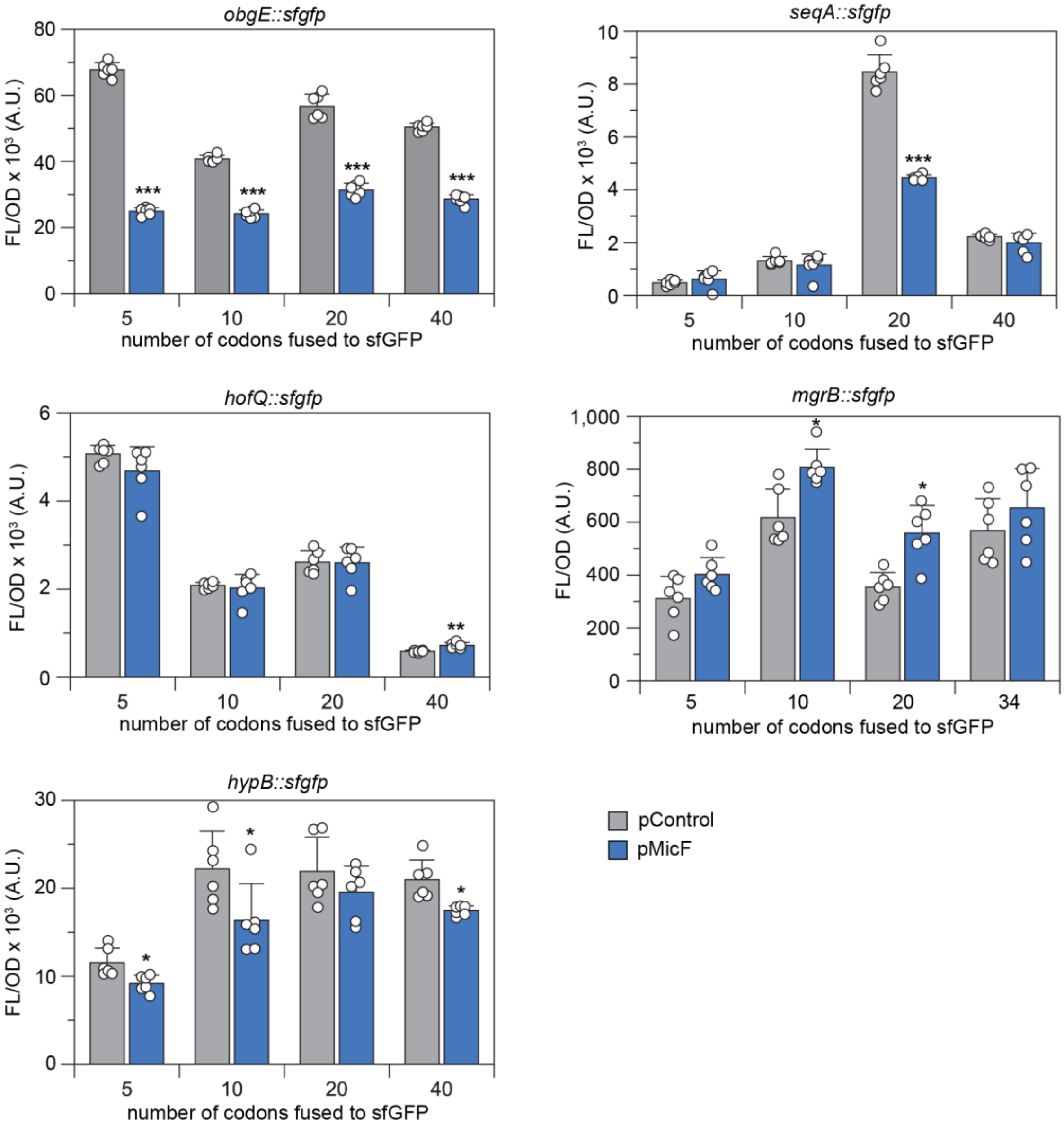


**Figure S1.** FL/$OD_{600}$ measurements for data in Figure 1. Bars show mean values and error bars represent standard deviation of six biological replicates shown as open circles. Two-tailed t-tests assuming unequal variance were used, and the significance is marked by asterisks above the bars indicating $p < 0.05$ (*), $p < 0.01$ (**), $p < 0.001$ (***).

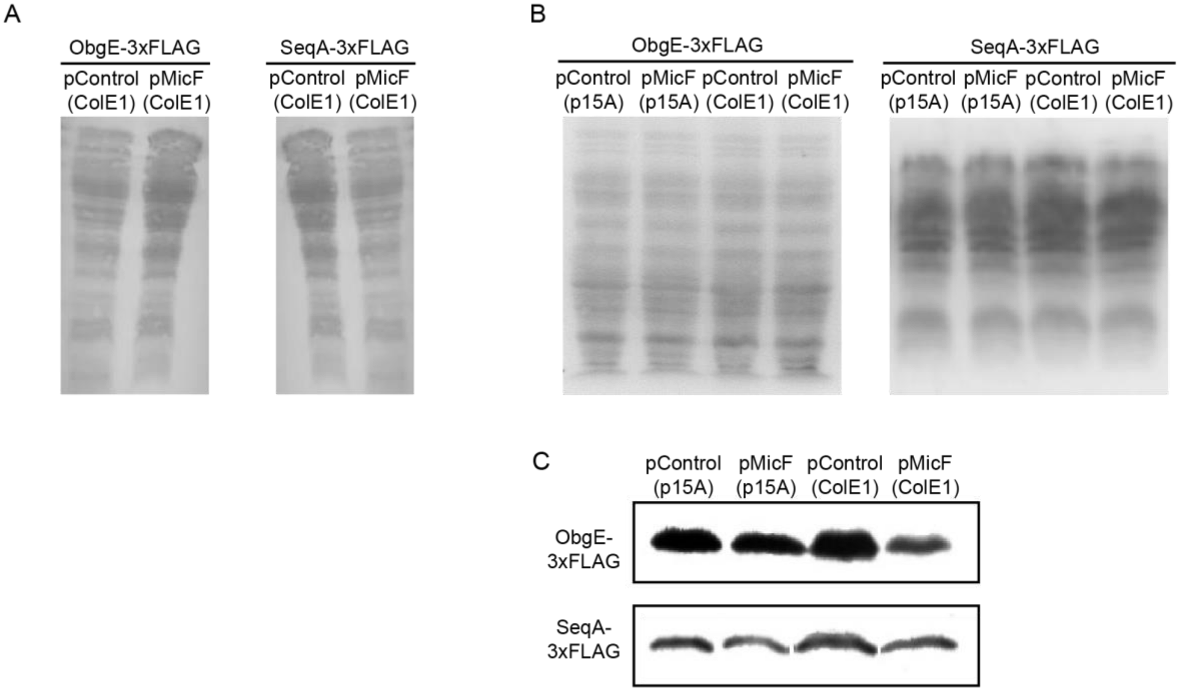


**Figure S2.** Controls and additional replicates for Western blots in Figure 2. A. Loading controls for Western blots in Figure 2 visualized via Ponceau S staining. B. Loading controls for Western blots in C visualized via Ponceau S staining. C. Additional Western blot analysis of ObgE-3xFLAG or SeqA-3xFLAG protein expressed in *E. coli* BW25113 $\Delta$*micF* transformed with either a MicF overexpression plasmid (pMicF) or control (pControl) on a high-copy vector (ColE1) or medium copy vector (p15A).

**Table S6**. Image J analysis of band intensities for ObgE-3xFLAG.

| | ColE1 – Figure 2 | | ColE1 – Figure SX | | p15A – Figure SX | |
|---|---|---|---|---|---|---|
| | pControl | pMicF | pControl | pMicF | pControl | pMicF |
| Loading control | 25309 | 28654 | 211102 | 221480 | 211097 | 221482 |
| Sample | 51585 | 16382 | 85454 | 11503 | 42587 | 36836 |
| Adjusted pControl/pMicF | 3.6 | | 7.8 | | 1.2 | |

Table SX. Image J analysis of band intensities for SeqA-3xFLAG

| | ColE1 – Figure 2 | | ColE1 – Figure SX | | p15A – Figure SX | |
|---|---|---|---|---|---|---|
| | pControl | pMicF | pControl | pMicF | pControl | pMicF |
| Loading control | 59100 | 26510 | 27243 | 24361 | 16750 | 17923 |
| Sample | 49854 | 9034 | 42373 | 28230 | 34051 | 23815 |
| Adjusted pControl/pMicF | 2.5 | | 1.3 | | 1.5 | |

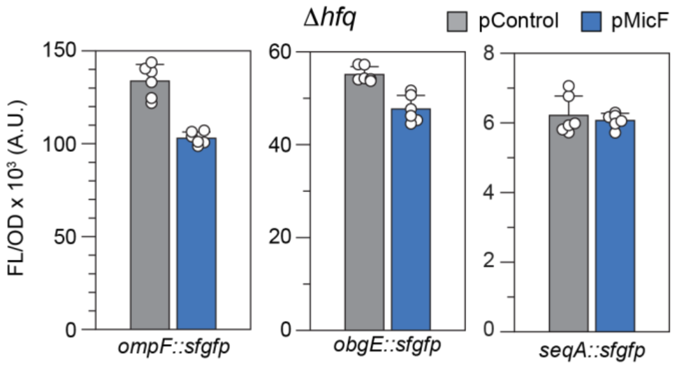


**Figure S3.** FL/$OD_{600}$ measurements for data in Figure 3. Bars show mean values and error bars represent standard deviation of six biological replicates shown as open circles.

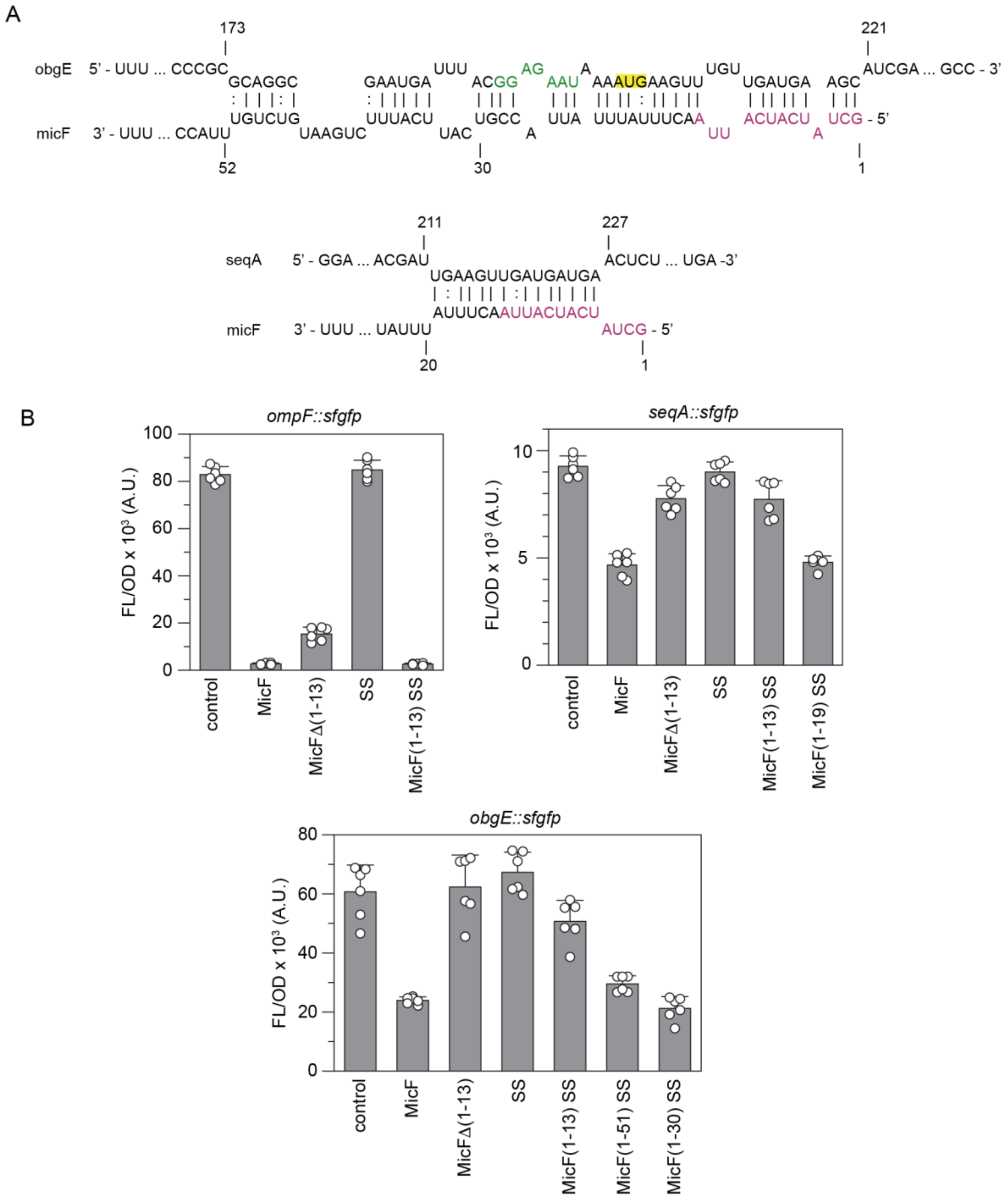


**Figure S4.** FL/$OD_{600}$ measurements for data in Figure 4. A. CopraRNA predicted MicF-mRNA interaction for *obgE* and *seqA.* Nucleotides in the predicted RBS of *obgE* are in green with the start codon highlighted in yellow. MicF's 13-nucleotide seed paring region is in magenta. B. Bars show mean values and error bars represent standard deviation of six biological replicates shown as open circles.

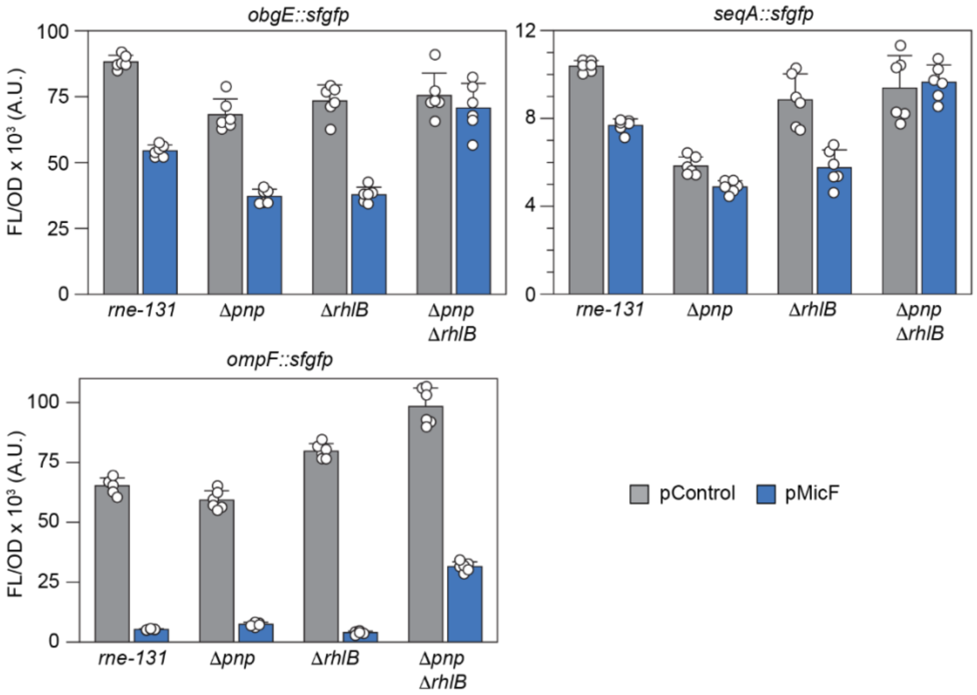


**Figure S5.** FL/$OD_{600}$ measurements for data in Figure 5. Bars show mean values and error bars represent standard deviation of six biological replicates shown as open circles.

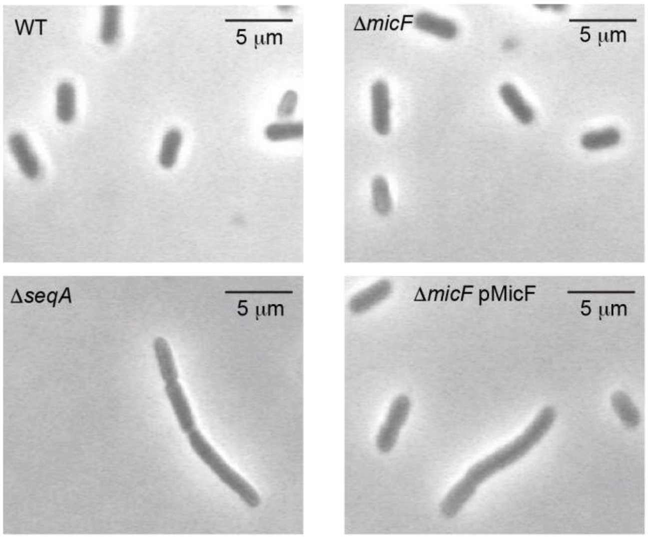


**Figure S6.** Additional observations of cell morphology (supplemental to Figure 6). *E. coli* BW25113 wild type, Δ*micF*, Δ*seqA*, and Δ*micF* complemented with a plasmid that overexpresses MicF (Δ*micF* pMicF) were grown to an $OD_{600}$ of 0.2 and visualized using phase contrast.

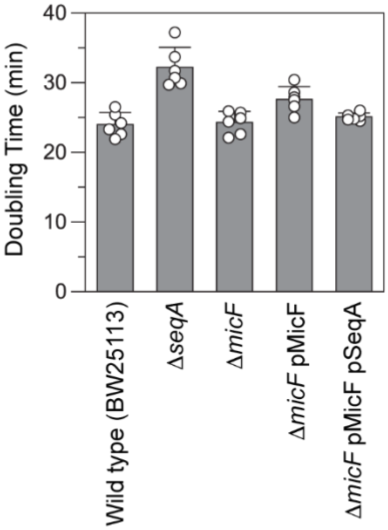


**Figure S7.** Doubling time measurements for data in Table 1. Bars show mean values and error bars represent standard deviation of six biological replicates shown as open circles.
